# Constitutive gene expression differs in three brain regions important for cognition in neophobic and non-neophobic house sparrows (*Passer domesticus*)

**DOI:** 10.1101/2021.07.06.451290

**Authors:** Christine R. Lattin, Tosha R. Kelly, Morgan W. Kelly, Kevin M. Johnson

## Abstract

Neophobia (aversion to new objects, food, and environments) is a personality trait that affects the ability of wildlife to adapt to new challenges and opportunities. Despite the ubiquity and importance of this trait, the molecular mechanisms underlying repeatable individual differences in neophobia in wild animals are poorly understood. We evaluated wild-caught house sparrows (*Passer domesticus*) for neophobia in the lab using novel object tests. We then selected the most and least neophobic individuals (n=3 of each) and extracted RNA from four brain regions involved in learning, memory, threat perception, and executive function: striatum, dorsomedial hippocampus, medial ventral arcopallium, and caudolateral nidopallium (NCL). Our analysis of differentially expressed genes (DEGs) used 11,889 gene regions annotated in the house sparrow reference genome for which we had an average of 25.7 million mapped reads/sample. PERMANOVA identified significant effects of brain region, phenotype (neophobic vs. non-neophobic), and a brain region by phenotype interaction. Comparing neophobic and non-neophobic birds revealed constitutive differences in DEGs in three of the four brain regions examined: hippocampus (12% of the transcriptome significantly differentially expressed), striatum (4%) and NCL (3%). DEGs included important known neuroendocrine mediators of learning, memory, executive function, and anxiety behavior, including serotonin receptor 5A, dopamine receptors 1, 2 and 5 (downregulated in neophobic birds), and estrogen receptor beta (upregulated in neophobic birds). These results suggest that some of the behavioral differences between phenotypes may be due to underlying gene expression differences in the brain. The large number of DEGs in neophobic and non-neophobic birds also implies that there are major differences in neural function between the two phenotypes that could affect a wide variety of behavioral traits beyond neophobia.

## Introduction

Neophobia (“fear of the new”) describes an animal’s reluctance to approach a novel object, try a new food, or explore an unfamiliar environment, behaviors that have been described in dozens of different animal species [1]. Neophobia is often repeatable within individuals [2, 3] and across contexts [4, 5], suggesting that it reflects an animal’s underlying exploratory temperament [6, 7]. A meta-analysis of personality traits in wild animals estimated the average heritability of exploration-avoidance behaviors (which includes novel object and novel environment tests) to be 0.58, suggesting a genetic basis to neophobia [8], and other studies have shown neophobia can be significantly influenced by parental identity [9] and early life environmental conditions [10].

A willingness to explore novelty may increase an individual’s likelihood of discovering new foods and nest sites, but it may also increase predation and disease risk [11–13]. Because novel urban and suburban environments are replacing natural environments on a global scale, neophobia is a personality trait with critical ecological and evolutionary relevance for wild populations [14]. Indeed, several studies have shown that neophobia affects animals’ ability to adapt to new challenges and opportunities [15–18], suggesting this personality trait is important in determining why some individuals, populations, and species are able to persist in human-altered landscapes whereas others are not.

Despite the ubiquity and importance of this personality trait, the neurobiological mechanisms underlying repeatable individual differences in neophobia behavior are not well understood in wild species. Next generation sequencing techniques have dramatically increased our ability to identify novel molecular mediators contributing to heritable and environmental causes of behavior by taking a data-driven approach [19–22]. Indeed, distinct patterns of neural gene expression can be associated with different behavioral types, as seen in species from honey bees [23] to stickleback fish [24]. Understanding more about the molecular mechanisms underlying neophobia may help us understand how this behavior develops, its genetic causes, and its fitness consequences – e.g., determining whether behavioral differences may be partly due to the presence of specific splice variants affecting the function of critical neural mediators of neophobia [25, 26].

In this study, we first screened a group of wild-caught house sparrows (*Passer domesticus*, n=15) for neophobia behavior in the lab using a set of novel objects placed on, in, or near the food dish. House sparrows are a highly successful invasive species displaying wide and repeatable individual variation in neophobia behavior in both the lab and the wild [27–30], have a sequenced genome [31, 32], and are a frequently used wild model system in endocrinology [33–35], immunology [36–38], and behavioral ecology [39–41]. This natural variation in neophobia makes house sparrows an excellent model to examine how individual variation in behavior may be linked to specific neurobiological differences. After neophobia screening, we selected a subset of the most and least neophobic individuals (n=3 of each) and extracted RNA from four candidate regions involved in learning, memory, threat perception, and executive function in birds: striatum, dorsomedial hippocampus, medial ventral arcopallium (AMV, previously referred to as the nucleus taenia of the amygdala), and caudolateral nidopallium (NCL; considered the avian “prefrontal cortex”) [42–47]. We created cDNA libraries and examined transcriptome differences in constitutive gene expression in these four brain regions. We had three main objectives for this project: 1) to determine whether overall patterns of constitutive gene expression differed in neophobic vs. non-neophobic individuals in our four regions of interest, 2) to identify differences in neurobiological pathways and processes in neophobic and non-neophobic birds, 3) to screen data from the first analysis to identify novel potential mediators of behavior that we or other researchers could examine in future studies.

## Methods

### Study subjects

House sparrows (n=15; 8 females, 7 males) were captured using mist nets at bird feeders in New Haven, CT, USA on 9 and 11 February 2018. Sparrows can be sexed using plumage features [48]; all animals were adults. In the lab, animals were singly housed with *ad libitum* access to mixed seeds, a vitamin-rich food supplement (Purina Lab Diet), grit, and water. Animals also had access to multiple perch types and a dish of sand for dustbathing. Animals were solo housed rather than group housed to avoid potential effects of social interactions on neophobia [30]. Day length in the lab corresponded to natural day length at the time of capture (10.5L:13.5D). Birds were allowed to habituate to laboratory conditions for 8 weeks before the start of experiments. Animals were collected under Connecticut state permit 1417011, and all procedures approved by the Yale University Animal Care and Use Committee under permit 2017-11648. We used approved methods for bird capture, transport, and husbandry as specified in the Ornithological Council’s Guidelines to the Use of Wild Birds in Research [49], and approved methods of euthanasia for avian species as specified in the 2020 American Veterinary Medical Association Guidelines for the Euthanasia of Animals.

### Neophobia protocol

Birds were fasted overnight and food dishes replaced in the morning 30 min after lights on with a novel object or the normal food dish alone (for control trials). Because birds do not eat in the lab after lights out (Supplemental Table S1), this only represents an additional 2 h of fasting at maximum for birds that do not feed during neophobia trials. After food dishes were replaced, behavior was video recorded for 1 h using web cameras (Logitech C615) connected to laptop computers to determine how long it took animals to approach and feed. Birds could not see each other during trials because of dividers placed between cages 24 h before the neophobia trials, although they could hear each other. Five different novel objects were used that either modified a normal silver food dish or were placed on, in, or near the food dish. These objects were: a normal silver food dish painted red on the outside (red dish), a red wrist coil keychain wrapped around the dish (ring), a blinking light hung above the dish and directed towards the front of the dish (light), a white plastic cover over part of the food dish (cover), and a green plastic egg placed on top of food in the middle of the dish (egg). These objects were used because they have been shown in another songbird species, the European starling (*Sturnus vulgaris*), to cause a significantly longer latency to approach compared to no object [50]. Some of these objects have also been shown to elicit neophobia in house sparrows [30]. Each bird was exposed to four of the five objects and four control trials (8 trials/bird, or 120 trials total). Video was lost from four trials (two control trials, two object trials) because of video camera malfunctioning, so final n=116 trials.

### Behavior data analysis

We investigated the effects of experimental condition (control or novel objects) and phenotype (neophobic, non-neophobic, or intermediate) on latency to feed with Cox proportional hazard models using the coxme package [51] in R Studio version 4.0.2 [52]. Using a survival analysis approach avoids having to create arbitrary threshold values when a subject does not perform the expected behavior during the allotted time period - i.e., giving subjects a time of 3600 s if they do not feed during a 60 min trial. All models included individual as a random effect. To ensure that the novel objects elicited neophobia, our first Cox proportional hazard model used experimental condition (object vs no object) as a fixed effect to estimate the overall effect of novel objects on latency to feed. We then ran a second model comparing each of the objects to control trials to estimate the effect of each object separately. Using average response times to feed during object trials, for our third and final model we split birds into three groups: strongly neophobic (n=3 females, 1 male), strongly non-neophobic (n=3 females, 1 male) and intermediate (n=3 females, 4 males). We then ran Cox proportional hazard models on novel object trials only to determine whether behavior in these groups was statistically different. This model included trial number as a fixed effect to examine possible habituation to novel object testing. We used log-rank post-hoc analyses in the survminer package [53] to compare average feeding times in the presence of novel objects among the three different phenotypes. We also examined repeatability in individual novel object responses using the ICC package, which calculates the intraclass coefficient [54]. For all models, we ensured that data met the assumptions of Cox models by testing the proportional hazards assumption using Schoenfeld residuals with the survival [55, 56] and survminer packages, and checking for influential observations by visualizing the deviance residuals using the survminer package. For all behavior analyses, *α* = 0.05, and because so few tests were used (3 total), we did not use multiple comparisons corrections.

### RNAseq tissue preparation and sample collection

Three weeks after the end of neophobia testing, the three most neophobic females and three least neophobic females were euthanized using an overdose of isoflurane anesthesia and brains rapidly removed and flash frozen in dry-ice cooled isopentane (Sigma Aldrich, St Louis, MO). We only used the females from the most and least neophobic groups to control for potential sex effects in gene expression; sex differences in neophobia behavior are not typically seen in this species [27, 29]. We stored brains at −80°C until sectioned coronally on a cryostat (Cryostar NX50, Thermo Fisher; −21°C) and mounted slices directly onto slides in two alternating series. The first series used 50 μm slices, dried overnight at 4°C and stained with thionin the following day. This first series of slides was used to help locate the brain regions of interest on the second series of slides. We sliced the second series at 200 μm, immediately transferred tissue to microscope slides on dry ice, and stored them at −80°C until extracting brain regions of interest. We sterilized the cryostat with a RNase/DNase removal reagent (DRNAse Free, Argos Technologies) followed by 95% ethanol, and replaced blades between subjects.

After confirming locations using the stained 50 μm series, we took brain tissue punches from four target brain regions: striatum mediale (striatum; did not include Area X), dorsomedial hippocampus [57], medial ventral arcopallium (AMV) [47], and caudolateral nidopallium (NCL) (Fig 1). We used the following punch sizes: striatum: 2 mm diameter (Fine Science Tools No. 18035-02, 11 G), hippocampus and NCL: 1 mm diameter (Fine Science Tools No. 18035-01, 15 G), and AMV: 0.5 mm diameter (Fine Science Tools No. 18035-50, 19 G). The striatum, hippocampus, and NCL are large brain regions. To ensure consistency in the relative position of the punches in the brain, we used other easily identified regions as landmarks: the start of the tractus quintofrontallis for the striatum, the start of the cerebellum for dorsomedial hippocampus, and NCL punches on the following slide from AMV. Brain regions were identified using published songbird brain atlases [58, 59] and house sparrow reference slides stained with thionin for DNA and Nissl substance and tyrosine hydroxylase to help locate NCL [46]. We combined one punch from each hemisphere (with the exception of AMV, in which case three smaller punches from each hemisphere were combined), in sterile, RNAse-free 1.6 mL centrifuge tubes submerged in dry ice and stored at −80°C until RNA extraction. We sterilized the punch tools in DPEC-treated water followed by D/RNAse Free (Argos Technologies) and 95% molecular-grade ethanol between subjects and brain regions.

**Figure 1.**
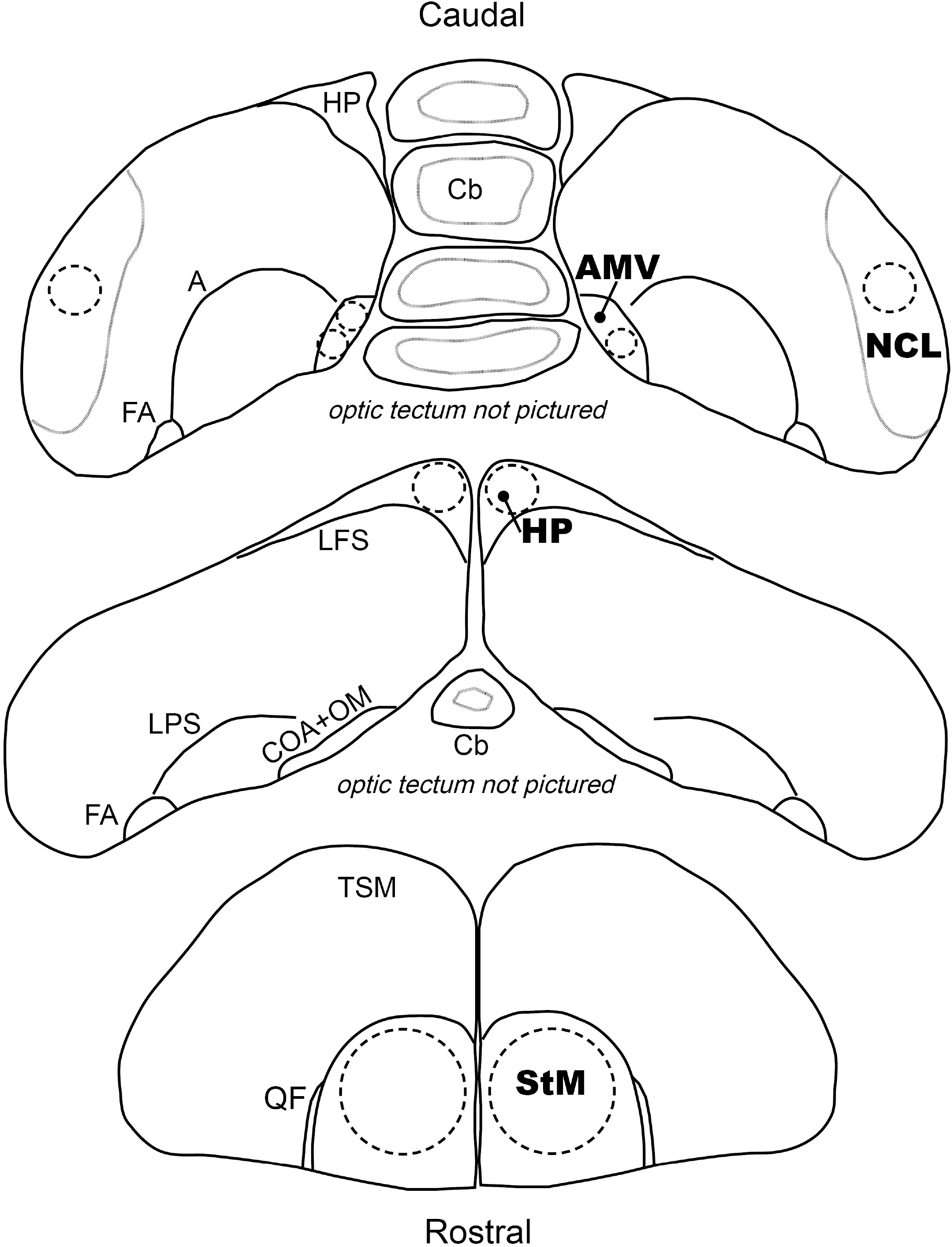
Locations of brain punches used for house sparrow RNAseq. Depiction of the approximate locations brain punches were taken from coronal 200 μm sections and the corresponding regions used as landmarks. **Top (caudal)**: Ventral medial arcopallium (AMV) samples consisted of three 11 G punches and caudolateral nidopallium (NCL) samples consisted of two 15 G punches. The NCL was sampled on the following section after AMV, but the regions are pictured on the same slice for simplicity. **Middle**: Dorsomedial hippocampus (HP) samples consisted of two 15 G punches. **Bottom** (rostral): Striatum (StM) samples consisted of two 18 G punches. **Abbreviations**: Cb = cerebellum, A = arcopallium, FA = tractus fronto-arcopallialis, LFS = lamina frontalis suprema, LPS = lamina pallio-subpallialis, COA = anterior commissure, OM = tractus occipito-mesencephalicus, TSM = tractus septopallio-mesencephalicus, QF = tractus quintofrontalis.

### RNA extraction and library preparation

We extracted RNA from brain tissue using the RNeasy® Lipid Tissue Mini Kit (QIAGEN; 1023539) and ran quality control on all samples using an Agilent 2100 Bioanalyzer system. The average RIN score for RNA samples was 8.5 (range 7.9 – 9.3). Extracted total RNA samples were sent to Novogene for library preparation and sequencing using 150 bp paired-end reads on a single lane of a NovaSeq 6000.

Sequencing of the mRNA libraries produced a total of 800 million 150 bp paired-end reads. All reads were trimmed of adapters and low quality bases using Trimmomatic (v.0.38) [60] with the following parameters (ILLUMINACLIP:TruSeq3-PE.fa:2:30:10 LEADING:3 TRAILING:3 SLIDINGWINDOW:4:15 MINLEN:36 or ILLUMINACLIP:TruSeq3-SE.fa:2:30:10 LEADING:3 TRAILING:3 SLIDINGWINDOW:4:15 MINLEN:36), and sequencing quality checked using the software FastQC (v.0.11.5) [61]. Trimmed reads were mapped to gene sequences annotated in the previously published genome for *P. domesticus* (GCA_001700915.1; [32]) using the two-pass mapping and transcriptome quantification modules in STAR (v.2.7.1; [62]). We used featureCounts (v.2.4.3) to extract read counts overlapping unique gene features, and measured differential expression in R (v.3.5.1) with the package edgeR (v.3.22.5) [63]. Sequence reads were filtered using a cutoff of 0.5 count per million in at least 6 samples, samples were normalized with post-filtering library sizes, and quasi-likelihood estimates of dispersion were calculated using the glmQLFit function. Global patterns in gene expression were analyzed using principal coordinate analysis (PCoA) using log-transformed reads generated with the cpm() function in edgeR with log=T and prior.count set to 1. These log-transformed reads were used to calculate dissimilarity indices using the R package vegan (v.2.5-5) and the pcoa function from the R package ape (v.5.3) [64, 65]. The influence of phenotype (P), brain region (BR), and their interactions with individual (I) were analyzed using the adonis2 permutational multivariate analysis of variance with the formula (P + BR + P*BR + BR*I) in vegan with 1e^6^ permutations.

Differential expression between treatments was tested using a combination of pairwise contrasts with the edgeR function glmQLFTest, as described by [66]. For these analyses, one model with no intercept was generated and grouped by brain region*phenotype. With this model, differential expression was measured between neophobic and non-neophobic individuals for each brain region independently. This approach allowed us to describe differences in constitutive levels of gene expression on a per-tissue basis, and in addition, because each individual had all 4 brain regions sequenced, we could also compare the resulting contrasts to identify shared and diverging responses between brain regions. This allowed us to better identify genomic markers that are specific to neophobia within and between each brain region. Genes identified as differentially expressed in these pairwise comparisons were tested for functional enrichment across all 3 major Gene Ontology classes (i.e., Biological Process (BP), Cellular Component (CC), and Molecular Function (MF)) and eu**k**aryotic **o**rthologous **g**roup (KOG) annotations with a Mann-Whitney U test in R using the ape package (v.5.2) [65] and code developed by [67]. For this analysis, the input for the Mann-Whitney U test was the negative log of the p-value for each gene multiplied by the direction of differential expression for that comparison, while the reference list was the complete list of genes included in the analysis. Finally, to capture a broader picture of the processes that were differentially regulated between neophobic and non-neophobic individuals, we tested for enrichment of KEGG pathways for each tissue type using the R package pathfindR! (v.1.4.2) [68].

## Results

### Behavior

Across all birds, the presence of a novel object at the food dish significantly increased the time to feed (β = −1.73, hazard ratio = 0.18 (confidence interval (CI): 0.11-0.29), z = −7.01, p < 0.0001) (raw behavior data and R code used for analysis are available as Supplemental Files 1-3). The latency to feed from a dish in the presence of any novel object was significantly longer than the control condition (control vs. keychain: p < 0.0001; control vs. red dish: p < 0.0001; control vs. light: p < 0.0001; control vs. egg: p = 0.012; control vs. cover: p = 0.0004). Considering only novel object trials, there was a significant difference in the latency to feed among birds classified as neophobic, non-neophobic, and intermediate (Fig 2; β = −0.84, hazard ratio = 0.43 (CI: 0.29-0.65), z = − 4.04, p = 0.0005). We did not detect an effect of trial number (β = 0.037, hazard ratio = 1.04 (CI: 0.92-1.18), z = 0.58, p = 0.56), suggesting that birds did not habituate to the testing procedure during novel object trials. Log-rank post-hoc analyses indicated that the neophobic birds were significantly different from both the non-neophobic birds (p = 0.00012) and intermediate birds (p = 0.00064) in their latency to feed in the presence of novel objects; however, intermediate and non-neophobic birds did not differ (p = 0.22). Including all three phenotypes, the intraclass correlation coefficient of the four individual novel object responses was 0.31 (CI: 0.06-0.62).

**Figure 2.**
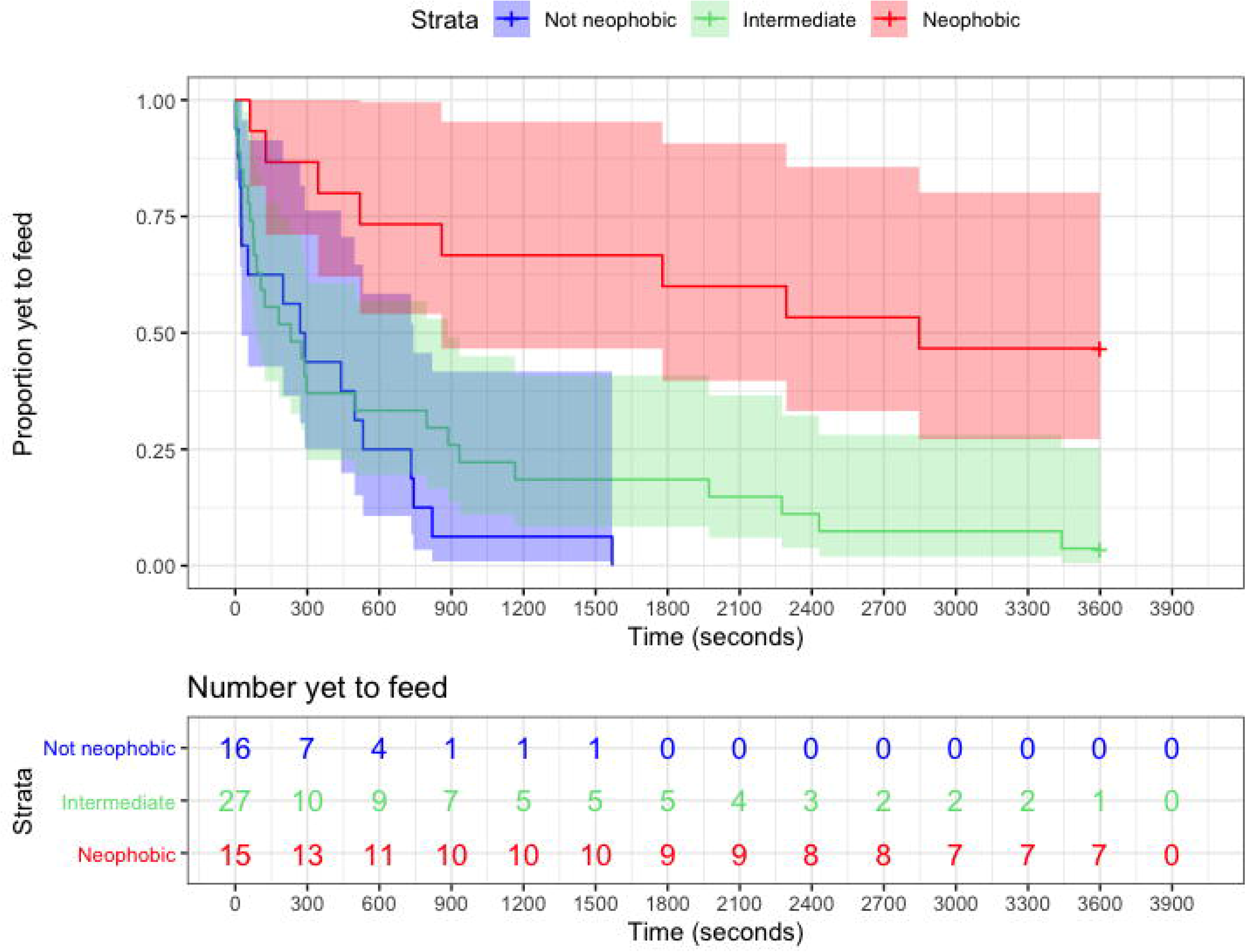
**Top**: Kaplan-Meier survival curves of house sparrow feeding likelihood in the presence of a novel object (four trials for each bird, except for one missing object trial for an intermediate bird where the video camera malfunctioned), split by neophobia phenotype (not neophobic n=4, intermediate n=7, neophobic n=4) and with 95% confidence intervals. **Bottom**: The risk table indicates the number of sparrows yet to feed from the dish in 300 s intervals. Both plot and table were created using the ‘survminer’ package in R Studio [53].

### RNAseq

Sequencing of the mRNA libraries produced a total of 800 million 150 bp paired-end reads (raw sequence data are being archived on the NCBI Single Read Archive (SRA) under accession SUB9422068). Read filtering for low quality scores left an average of 32.2 million reads per sample (range: 24.5 - 42.6 million). Mapping of these reads to the previously published house sparrow reference genome [31] resulted in an average of 80% unique mapping rate (range: 75% - 83%). For our analysis, we focused the analysis on the 13,193 gene regions annotated in the reference genome (GCA_001700915.1; [32]). The gene set was filtered to remove features that did not have at least 0.5 counts-per-million reads in 25% samples. The final differential expression analysis was run on these 11,889 genes for which we had an average of 25.7 million mapped reads per sample (range: 19.2 – 34.3 million mapped reads per sample). PERMANOVA results identified significant effects of brain region, phenotype (neophobic vs. non-neophobic), and a brain region by phenotype interaction, but no effect of individual identity on gene expression (Fig 3). Below we briefly describe the observed transcriptomic signatures of neophobic behavior for each brain region. For all analyses, significantly differentially expressed genes (DEGs) are those with a logFold change greater than 1 or less than −1 and a Benjamini-Hochberg corrected false discovery rate (FDR) less than or equal to 0.05.

**Figure 3.**
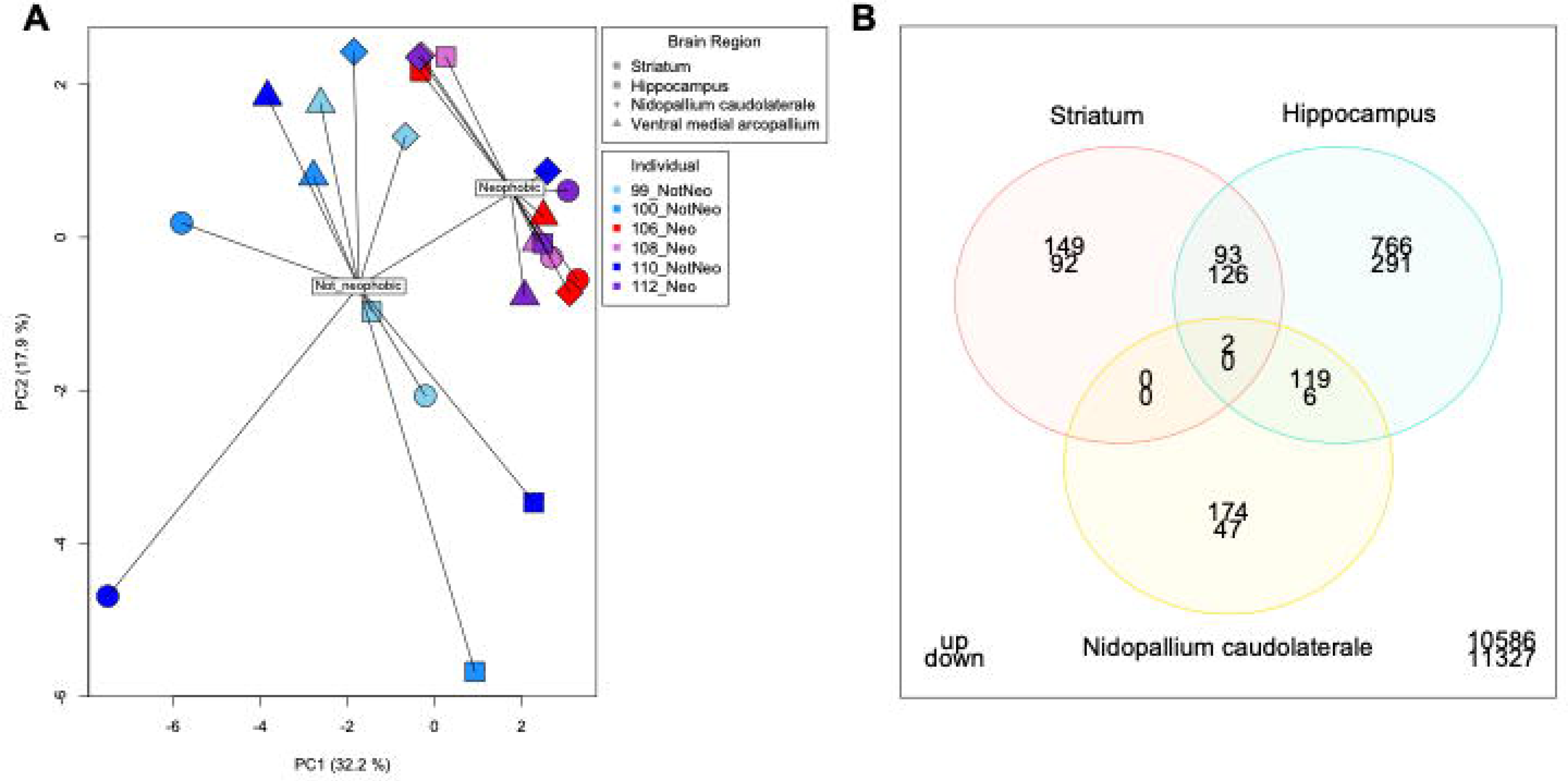
A) Principal coordinate analysis (PCoA) spider plot of gene expression, split by brain region and phenotype. Each brain region is represented by a different shape, and phenotypes are represented by colors (blue shades: not neophobic or “NotNeo”, n=3 and red shades: neophobic or “Neo”, n=3). Results from permutational multivariate analysis of variance (PERMANOVA) are shown. B) Venn diagram of genes differentially expressed between neophobic and not neophobic individuals, highlighting minimal overlap among brain regions in the identities of differentially expressed genes.

### 1. Hippocampus

Differential gene expression analysis for hippocampus samples identified 1,403 DEGs (12% of the measured transcriptome) with 980 genes upregulated and 423 genes downregulated in neophobic birds relative to non-neophobic birds (for this and other regions, see Supplemental File 4 for full list). Genes showing the strongest downregulation among neophobic individuals included a cytosolic phospholipase A2 gene member (PLA2G4E; logFC = −9.7, adj. p value = 0.026); membrane metallo-endopeptidase or neprilysin, a zinc-dependent metalloprotease (MME; logFC = −6.7, adj. p value = 0.009); probable vesicular acetylcholine transporter-A (slc18a3a; logFC = −5.8, adj. p value = 0.007); and protachykinin-1 (TAC1; logFC = −5.2, adj. p value = 0.004). In addition to these, there were 3 dopamine receptors that were significantly downregulated (DRD1; logFC = −2.9, adj. p value = 0.004 | DRD2; logFC = −6.3, adj. p value = 0.007 | DRD5; logFC = −3.4, adj. p value = 0.008). Genes showing the strongest upregulation among neophobic individuals included the estrogen receptor beta gene (ERß; logFC =9.6, adj. p value =0.012; an odd-skipped-related 1 gene (osr1; logFC =7.9, adj. p value = 0.038); a transthyretin gene (TTR; logFC = 7.6, adj. p value =0.02); and a gene coding for lipocalin (Lipocalin; logFC = 6.9, adj. p value = 0.044). The Fisher’s exact test with upregulated genes found 46 enriched Molecular Function (MF) ontologies, 189 Biological Process (BP) ontologies, and 62 Cellular Component (CC) ontologies. This same test for enrichment with downregulated genes only identified 9 ontologies, all of which were CC terms (for this and other regions, see Supplemental File 5 for full list).

Measuring global differences in expression with a eukaryotic orthologous group (KOG) enrichment analysis identified 3 enriched KOG terms with decreased expression of genes involved in translation, energy production, and metabolism in neophobic birds relative to non-neophobic birds (Fig 4). We also observed 5 enriched KOG terms among the upregulated genes that were involved in cellular structure, signal transduction, and posttranslational modifications. Together, the two ontology-based analyses found that the majority of enriched terms were found among upregulated transcripts in neophobic birds, and were broadly distributed across structural, signaling, and metabolic processes (for this and other regions, see Supplemental File 6 for full list).

**Figure 4.**
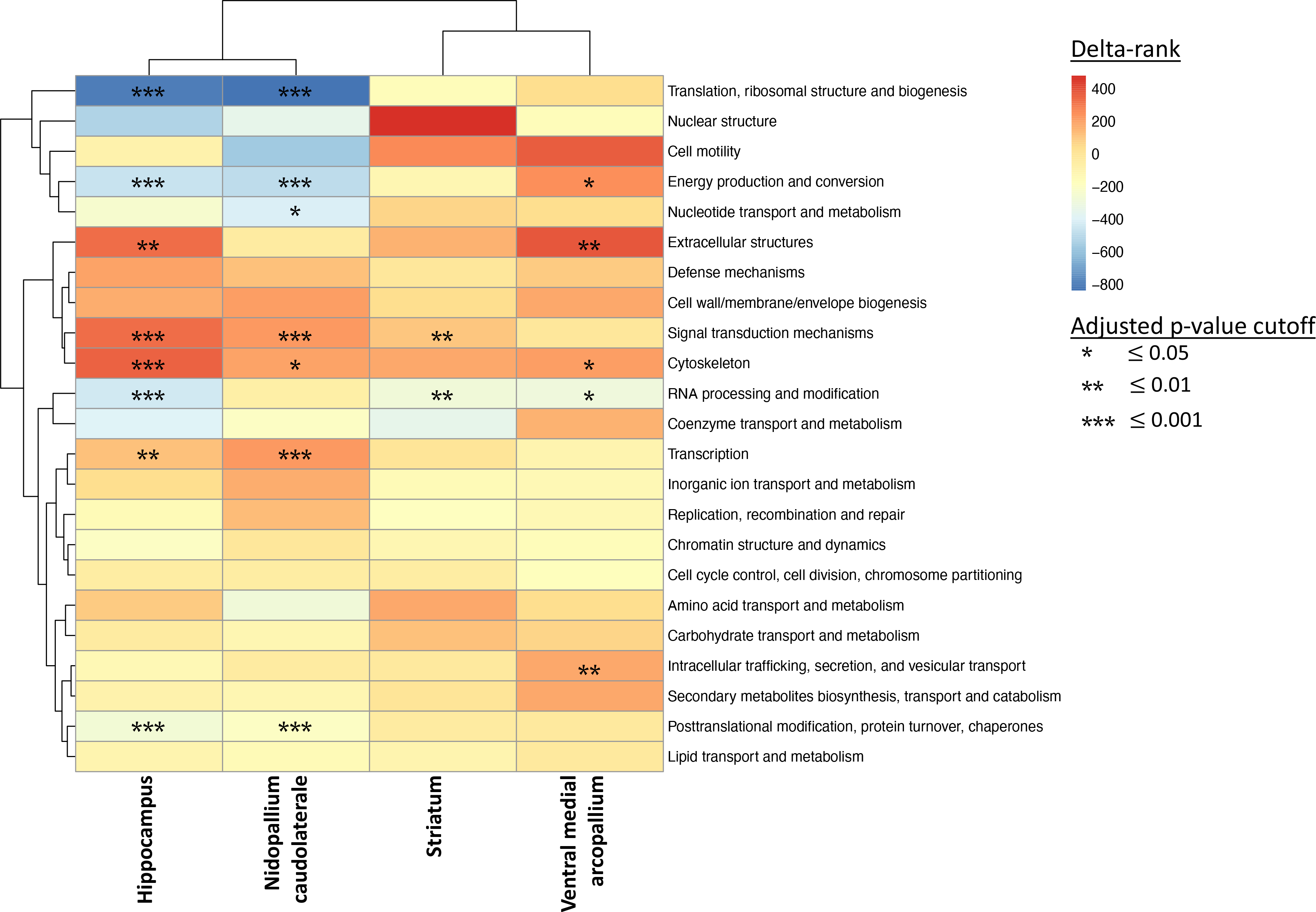
Enriched eukaryotic orthologous group (KOG) terms in the house sparrow transcriptome across four brain regions. Positive delta-ranks (red) are associated with upregulation in neophobic birds relative to non-neophobic birds, and significance is based on Benjamini-Hochberg adjusted p-values (FDR).

### 2. Striatum

Differential gene expression analysis for striatum samples identified 462 DEGs between phenotypes, with 244 upregulated genes and 218 downregulated genes in neophobic birds relative to non-neophobic birds. Genes with the strongest downregulation among neophobic individuals included a metallophosphoesterase 1 gene (MPPE1; logFC = −11.7, adj. p value = 0.027); a transmembrane protein 61 gene (TMEM61; logFC = −8.1, adj. p value = 0.014); and a GRB2-associated-binding protein 2 (GAB2; logFC = −6.8; adj. p value = 0.035). The upregulated genes in this comparison included a protein N-terminal asparagine amidohydrolase (NTAN1; logFC = 9.1, adj. p value = 0.016); multiple NADH dehydrogenases (NDUFV1; logC = 5.3, adj. p value = 0.03 | NDUFB3; logFC = 5.2, adj. p value = 0.02 | NDUFA6; logFC = 2.12, adj. p value = 0.04); and superoxide dismutase (SOD; logFC = 3.8, adj. p value = 0.025). Examining functional enrichment among upregulated genes with the Fisher’s Exact Test found no significant enrichment for MF terms, 1 for BP terms, and 7 for CC terms. There was also a small number of enriched terms among downregulated transcripts, with 10 enriched terms identified. These results are mirrored in the KOG analysis, with 1 enriched term among upregulated transcripts and 1 enriched term among downregulated transcripts (Fig 3). These enriched terms reveal decreased expression of genes associated with RNA processing and increased expression among signal transduction pathways.

### 3. Caudolateral nidopallium (NCL)

Differential gene expression analysis for the NCL samples found 348 DEGs between phenotypes, with 295 upregulated and 53 downregulated genes in neophobic birds relative to non-neophobic birds. The strongest downregulated genes were a voltage-gated potassium channel (KCNG4; logFC = −5.6, adj. p value = 0.032); serotonin receptor 5A (HTR5A; logFC = −3.5, adj. p value = 0.044); and a transmembrane protein potentially associated with endocytosis (CHODL; logFC = −3.2, adj. p value = 0.043). The upregulated genes were found to include a nuclear envelope protein (SYNE2; logFC = 5.0, adj. p value = 0.013); a gene important for active DNA demethylation (TET1; logFC = 3.1, adj. p value = 0.013); and a calcium channel normally associated with cardiac muscle (RYR2; logFC = 2.7, adj. p value = 0.019). Gene ontology analysis using the Fisher’s Exact Test found strong levels of enrichment for the 295 upregulated genes with 34 enriched ontologies associated with MF, 111 enriched BP ontologies, and 52 enriched CC ontologies. In addition, KOG enrichment analysis found 7 enriched KO terms all of which were shared with the enrichment observed in samples from dorsomedial hippocampus (Fig 3).

We explored potential drivers of these shared responses by comparing the genes that were differentially expressed in both NCL and hippocampus samples. This comparison identified 129 genes differentially expressed in both brain regions. Of these, 121 were found to be upregulated in both tissues, while only 6 genes were found to be downregulated in both tissue types. The remaining 2 genes were differentially expressed in both tissue types but had opposing expression patterns, with higher expression observed in hippocampus samples. In both brain regions, we also observed enrichment for transcription, cytoskeleton, and signal transduction mechanisms among upregulated genes (Fig 3). The genes that were shared included SYNE2, TET, and five isoforms of a DST gene, all of which were upregulated in neophobic individuals. KEGG pathway analysis also identified similar patterns in pathway enrichment between the two brain regions associated with multiple signaling pathways, including Notch signaling, mTOR signaling, and insulin signaling.

### 4. Ventral medial arcopallium (AMV)

Differential gene expression analysis for AMV samples found no DEGs between neophobic and non-neophobic individuals. This lack of differential gene expression could be due to differences in brain punches used for this region; because of small region size, brain punches centered on this region also contained some of the surrounding regions. Despite this lack of significantly DEGs, we still explored functional enrichment using the Mann-Whitney U-test, which did identify significant enrichment for increased expression of oxidation-reduction processes and autophagy in neophobic birds relative to non-neophobic birds. There was also a decrease in expression for genes associated with transcription regulation, chromatin organization, and mRNA processing in neophobic birds. KOG enrichment analysis also identified significant enrichment for intracellular trafficking, extracellular structures, and energy production.

## Discussion

Similar to previous studies, we found large individual variation in neophobia in wild-caught house sparrows [27–29]. Based on average responses to novel objects, we split sparrows into highly neophobic, highly non-neophobic, and intermediate groups, and sequenced total mRNA libraries from four brain regions of three of the most and least neophobic individuals. Overall, we found that the three highly neophobic individuals we sequenced had very different patterns of constitutive gene expression in the brain compared to the three non-neophobic individuals. This project adds to a growing body of work showing distinct patterns of gene expression in the brain associated with different behavioral types [23, 24, 69–71].

Gene expression patterns in the dorsomedial hippocampus were especially distinct, where 12% of the transcriptome was differentially expressed in neophobic birds compared to non-neophobic birds, but also in the striatum and NCL, where 4% and 3% of genes were differentially expressed, respectively. These results suggest that these regions all play critical direct or indirect role in deciding whether or not to approach an unfamiliar object, and may therefore be important in evaluating potential threats and resources (exploratory behavior). Although studies have examined shared neural substrates for social and appetitive behavior across vertebrates [72], much less is known about possible conserved networks of brain regions involved in mediating *aversive* behavior. And while neural circuits involved in song learning, reproduction, and spatial learning have been particularly well-studied in songbirds [73–77], there are still many regions that are poorly understood in the avian brain. Therefore, this study provides essential data about the role of different brain regions in behavior that is often lacking outside of mammalian model systems. The large number of differentially expressed genes in the hippocampus in particular suggests this region merits a closer look as a potential driver of variation in personality traits like neophobia. However, one important limitation of this study was that only females were used, and future work should confirm that these patterns hold true for male sparrows as well.

Interestingly, despite previous work showing the involvement of the AMV (previously called nucleus taenia of the amygdala) in decision making and emotional responses involved in fear and anxiety [44, 45] and even to novelty specifically [78], there were no significant differences in constitutive gene expression between neophobic and non-neophobic animals in this brain region. While this may be due to methodological reasons (AMV is a smaller region, so our punches may have included more non-target tissue), this also suggests that differences in behavior between neophobic and non-neophobic birds are not driven by differences in the AMV. Indeed, submitted work from our lab examining immediate early gene activity in neophobic and non-neophobic birds demonstrates that both phenotypes show a similar increase in neuronal activity in the AMV response to novel objects compared to non-object controls [79].

Intriguingly, some of the most highly differentially expressed genes between neophobic and non-neophobic individuals include important known neuroendocrine mediators of learning, memory, executive function, and anxiety behavior, including serotonin receptor 5A, dopamine receptors 1, 2 and 5, and estrogen receptor beta [80–84]. Behavioral variation has been associated with differential receptor density and gene expression in specific neuromodulatory systems in several species. This includes differences in pallial glutamate receptors in wild finches with divergent problem-solving strategies [85], in forebrain serotonin receptors in salmon with different emergence times from spawning nests [86], and in whole brain benzodiazepine receptors in lizards with different behavioral responses to simulated predators [87]. Although genes with the highest fold change do not necessarily have the highest biological significance, these receptors are strong candidates for future work. Dopamine receptor 2 specifically has already been linked to personality traits such as boldness and novelty seeking in other species [88–90].

Surprisingly, neophobic birds in our study showed no evidence for differential expression of the dopamine receptor 4 (DRD4) gene in any of the four brain regions we examined. In fact, this gene was not consistently expressed in enough birds to be included in our analysis. DRD4 is one of the most commonly implicated candidate genes underlying variation in neophobic behaviors in birds, with polymorphisms in this gene linked to response to novelty in flycatchers [91], flight distance in dunnocks [92], wariness in swans [93], and invasion success in weavers [94]. Although we observed no differences in the expression of DRD4 in neophobic and non-neophobic birds, we did observe differential expression of dopamine receptors DRD1, DRD2 and DRD5, suggesting neophobic behaviors in different species may evolve through convergent changes targeting different genes in the same neuroendocrine systems. Similarly, we saw no differential expression in the serotonin transporter (SERT) gene, which has also been implicated in neophobic behaviors in several species [92, 95, 96]. Similar to DRD4, SERT was dropped from analysis because it was not consistently expressed. However, we did observe differential expression of the HTR5A (serotonin receptor 5A) gene, once again pointing to the possibility of convergent evolution through changes to different genes in the same neurotransmitter system. Alternatively, it is possible that in the other studies implicating DRD4 and SERT in neophobic behaviors, the observed polymorphisms are linked to protein coding, rather than regulatory changes, so those studies would also not have observed constitutive differences had they measured expression in these genes. While DRD4 and SERT are commonly implicated in variation in neophobic behavior, other studies have failed to find evidence for the involvement of one or both of these genes in neophobic behaviors [93, 96, 97], and in cases where these two genes were the only ones considered, some of these studies were unable to identify other candidates. Our findings highlight the utility of a comparative transcriptomic approach when attempting to understand behavioral variation in natural populations: by taking a global view of neurophysiological differences among individuals we were able to identify candidate genes not previously implicated in neophobia.

Across all four brain regions, KEGG pathway analysis showed a strong functional similarity between genes differentially expressed in neophobic birds in the hippocampus and in the NCL. This was somewhat unexpected because these two regions are not known to be directly connected in the avian brain [57, 98]; instead, the dorsomedial hippocampus reciprocally connects with the posterior pallial amygdala, which receives projections from the NCL [99]. Interestingly, in neophobic birds, translation, post-translation modification, and energy production and conversion pathways were underexpressed, while transcription-related genes and signal transduction pathways were overexpressed in these two regions. This suggests that in neophobic birds, transcription is increased but translation is decreased. This could affect behavioral plasticity in ways that remain to be explored with potential implications for small non-coding RNA and RNA-mediated processes being used more often in neophobic individuals. This may also relate to the increased expression of genes involved in post-translational modifications, protein turnover, and chaperone genes in NCL and hippocampus in neophobic birds. Further, in all regions but the AMV, genes associated with the signal transduction mechanisms pathway were overexpressed in neophobic birds relative to non-neophobic birds.

Importantly, as in many transcriptomics studies, the data presented here represent a single snapshot of gene expression. Gene expression in avian brains is highly dynamic, and large numbers of genes may be differentially expressed due to changes in a few ‘master regulators’ of gene expression [100]. As a result, we do not know which of the many genes that were differentially expressed in neophobic birds are actually causing these behavioral differences. Future work could examine differential expression in neophobic birds through time to help clarify differences in regulatory networks among behavioral phenotypes [101, 102].

In summary, we found that the brains of animals with different personality types differed in constitutive gene expression in three of the four brain regions we examined. Because these differences were present in the absence of novel stimuli, the large number of DEGs in neophobic and non-neophobic birds implies that there are major differences in neural function between the two phenotypes that could affect a wide variety of behavioral traits beyond neophobia, potentially leading to the existence of behavioral syndromes [103]. Because differences in gene expression do not necessarily mean differences in protein expression [104], future studies should use techniques like immunohistochemistry and Western blots to examine whether particular mediators in fact differ in protein expression in neophobic and non-neophobic birds. The cause of differences between neophobic and non-neophobic individuals is still unknown, but could include genetic variation [e.g., 105], epigenetics [e.g., 106], or environmental conditions during development and adulthood [e.g., 107, 108]. Understanding the neurobiological basis for different animal temperaments has important implications for ecology and evolutionary biology because it can affect macro-level processes such as species’ distributions and their ability to respond to environmental changes and exploit novel resources.

## Acknowledgements

Thanks to G. Cameron, R. Prum, and M. and P. Wolter for help acquiring sparrows, K. Elwell, W. Daniels and D. Torres for animal care, and G. Kim and C. Lu for assistance with neophobia trials. The authors also appreciate facilities and equipment support from R. Carson.

