## Supplemental Table S1 for "Constitutive gene expression differs in three brain regions important for cognition in neophobic and non-neophobic house sparrows (*Passer domesticus*)"

**Table S1.** House sparrows do not feed in the lab when lights are off overnight. Food dishes were weighed immediately after the lights in the bird room turned off in the evening. Dishes were re-weighed just before the lights turned on in the morning. Each bird was individually housed in a cage with a single food dish. Any differences in food cup mass between night and morning appear attributable to normal variation in mass measurements from the scale used (Mettler Toledo TLE3002E).

| **Sparrow ID** | **Night food cup mass** | **Morning food cup mass** | **Mass difference** |
| --- | --- | --- | --- |
| 87 | 46.65 | 46.65 | 0 |
| 88 | 46.07 | 46.08 | -0.01 |
| 91 | 46.37 | 46.38 | -0.01 |
| 95 | 46.64 | 46.60 | 0.04 |
| 94 | 47.30 | 47.31 | -0.01 |
| 106 | 45.43 | 45.41 | 0.02 |
| 112 | 47.65 | 47.65 | 0.00 |
| 107 | 46.42 | 46.43 | -0.01 |
| 98 | 47.08 | 47.08 | 0 |
| 110 | 46.39 | 46.39 | 0 |
| 116 | 45.24 | 46.23 | -0.99 |
| 144 | 46.96 | 46.96 | 0 |
| *Average:* | *46.52* | *46.59* | *-0.08* |
